# Cryogenic OrbiSIMS Localizes Semi-Volatile Molecules in Biological Tissues

**DOI:** 10.1101/862391

**Authors:** Clare L. Newell, Jean-Luc Vorng, James I. MacRae, Ian S. Gilmore, Alex P. Gould

## Abstract

OrbiSIMS is a recently developed instrument for label-free imaging of chemicals with micron spatial resolution and high mass resolution. Here we report a cryogenic workflow for OrbiSIMS (Cryo-OrbiSIMS) that improves chemical detection of lipids and other biomolecules in tissues. Cryo-OrbiSIMS decreases ion-beam induced fragmentation, allowing large molecules such as triglycerides to be more reliably identified. It also increases chemical coverage to include biomolecules with intermediate or high vapor pressures, such as free fatty acids and semi-volatile organic compounds (SVOCs). We find that Cryo-OrbiSIMS reveals the hitherto unknown localization patterns of SVOCs with high spatial and chemical resolution in diverse plant, animal and human tissues. We also show that Cryo-OrbiSIMS can be combined with genetic analysis to identify enzymes regulating SVOC metabolism. Cryo-OrbiSIMS is applicable to high resolution imaging of a wide variety of non-volatile and semi-volatile molecules across many areas of biomedicine.

## Main

Numerous semi-volatile organic compounds (SVOCs) such as aldehydes, esters and hydrocarbons are synthesized by plants, humans and other animals. SVOCs perform many different biological functions, with several acting as signals that convey pheromonal communication between individuals of the same or other species (Liberles, 2014; Xu and Turlings, 2018; Yew and Chung, 2015). Mass spectrometry of headspace vapors, impregnated fibers or solvent extracts has traditionally been used to analyze SVOCs produced by biological tissues (Lucattini et al., 2018). These methods are very sensitive but do not localize SVOCs in tissues with high spatial resolution. Chemical imaging of hydrocarbons is possible using matrix assisted laser desorption ionization (MALDI) and ultraviolet laser desorption ionization (UV-LDI) methods (Vrkoslav et al., 2010; Yew et al., 2011). As both techniques are limited by laser spot size, it is desirable to develop higher spatial resolution methods such as secondary ion mass spectrometry (SIMS). SVOCs are, however, challenging to analyze with SIMS as they are volatile under the ultra-high vacuum required by these instruments, and therefore cannot be imaged. To overcome this limitation, cryogenic temperatures can be used to lower the vapor pressures of SVOCs. Cryogenic workflows that prevent the ice sublimation of frozen-hydrated samples have been developed for SIMS instruments with a time-of-flight detector (ToF-SIMS) (Dickinson et al., 2006; Kuroda et al., 2013). ToF detectors have excellent sensitivity but, in many cases, they do not have sufficient mass resolution to assign peaks with high confidence.

The OrbiSIMS is a new hybrid SIMS instrument that allows chemical imaging with micron spatial resolution via Orbitrap and ToF detectors (Passarelli et al., 2017). The Orbitrap analyzer provides high mass resolution (>240,000 at *m/z* 200) with <2 ppm mass accuracy, enabling peaks to be assigned with a greater degree of confidence than is possible with a ToF detector, typically with mass resolution ~18,000 at *m/z* 200. We therefore set out to develop a cryogenic workflow for OrbiSIMS that would allow SVOCs to be imaged with high chemical and spatial resolution (Materials and Methods). As a test case, we chose hydrocarbons because they are biologically important SVOCs but many are volatile under high vacuum at ambient temperature (**Supplementary Fig. 1**). Hydrocarbons are extensively fragmented by many commonly used ion sources (Roussis et al., 1998), making it difficult to identify the parent ions. As hydrocarbon adducts have not previously been reported in SIMS, we first analyzed their chemical composition via Cryo-OrbiSIMS in positive polarity spectrometry mode - using either the 20 keV argon gas cluster ion beam (GCIB) or the 60 keV Bi_3_^++^ liquid metal ion gun (LMIG). Analysis of 9(Z)-tricosene, an insect pheromone hydrocarbon (Blomquist and Bagneres, 2010), showed that the 20 keV Ar_3500_^+^ GCIB tends to generate [M+N]^+^ ions, which may be formed from direct interactions with residual nitrogen gas in the instrument or via multiple hydrogen loss from [M+NH_4_]^+^ adducts (**Supplementary Fig. 2**). In contrast, the 60 keV Bi_3_^++^ LMIG primarily yields [M-2H]^+^ ions, as previously reported for GC-MS(McLafferty and Turecek, 1993), but leads to greater fragmentation of tricosene. GCIB analysis of an American Society for Testing and Material (ASTM) reference gas oil, a mix including a range of C10:0 to C52:0 alkanes, also showed abundant [M+N]^+^ ions (**Supplementary Fig. 3**). Hydrocarbon peak assignments were validated by comparing MS and MS/MS spectra from the OrbiSIMS Orbitrap with those from direct infusion HESI-Orbitrap and GC-MS (**Supplementary Fig. 4**). These findings show that a cryogenic OrbiSIMS workflow reliably detects hydrocarbons and they identify the major adducts formed.

Chemical imaging of human latent fingerprints has emerged as an important area of forensic and biomedical science. Several imaging technologies including ToF-SIMS have previously been used to localize non-volatile drugs, poisons, antibiotics and other molecules to the ridges and sweat pores of fingerprints (Cai et al., 2017; Hazarika and Russell, 2012; Ifa et al., 2008). Comparison of cryo- and ambient OrbiSIMS Orbitrap spectra showed the presence of many peaks at −110°C that are not observed at 30°C (**Fig. 1**). Orbitrap peak assignments indicate that many such peaks correspond to fatty acids and their derivatives as well as triacylglycerides (TAGs) and phosphoinositol-ceramides (PI-Cer) (**Supplementary Table 1**). Fatty acids are known to be abundant in latent fingerprints but have not been robustly detected in previous fingerprint studies using SIMS at ambient temperature, presumably because they are volatile under these high-vacuum conditions (**Supplementary Fig. 1**). Consistent with this, signals from C16:1 and C16:0 were reproducibly detected using OrbiSIMS in the cryogenic but not in the ambient mode. It is therefore surprising that previous studies using SIMS at ambient temperature have reported free fatty acids in other tissues (Brulet et al., 2010; Richter et al., 2007). One possible explanation is that the appearance of free fatty acids at ambient temperature results from beam-induced fragmentation of TAGs and/or other high molecular weight lipids that are not themselves volatile under ultra-high vacuum. In line with this, Ar_3500_^+^-Orbitrap OrbiSIMS analysis of latent fingerprints at ambient, compared to cryogenic, temperature shows decreased TAG and increased diacylglyceride signals (**Supplementary Table 1**). We conclude that cryogenic sample analysis improves OrbiSIMS detection of biologically important lipids in their intact state.

**Figure 1.**
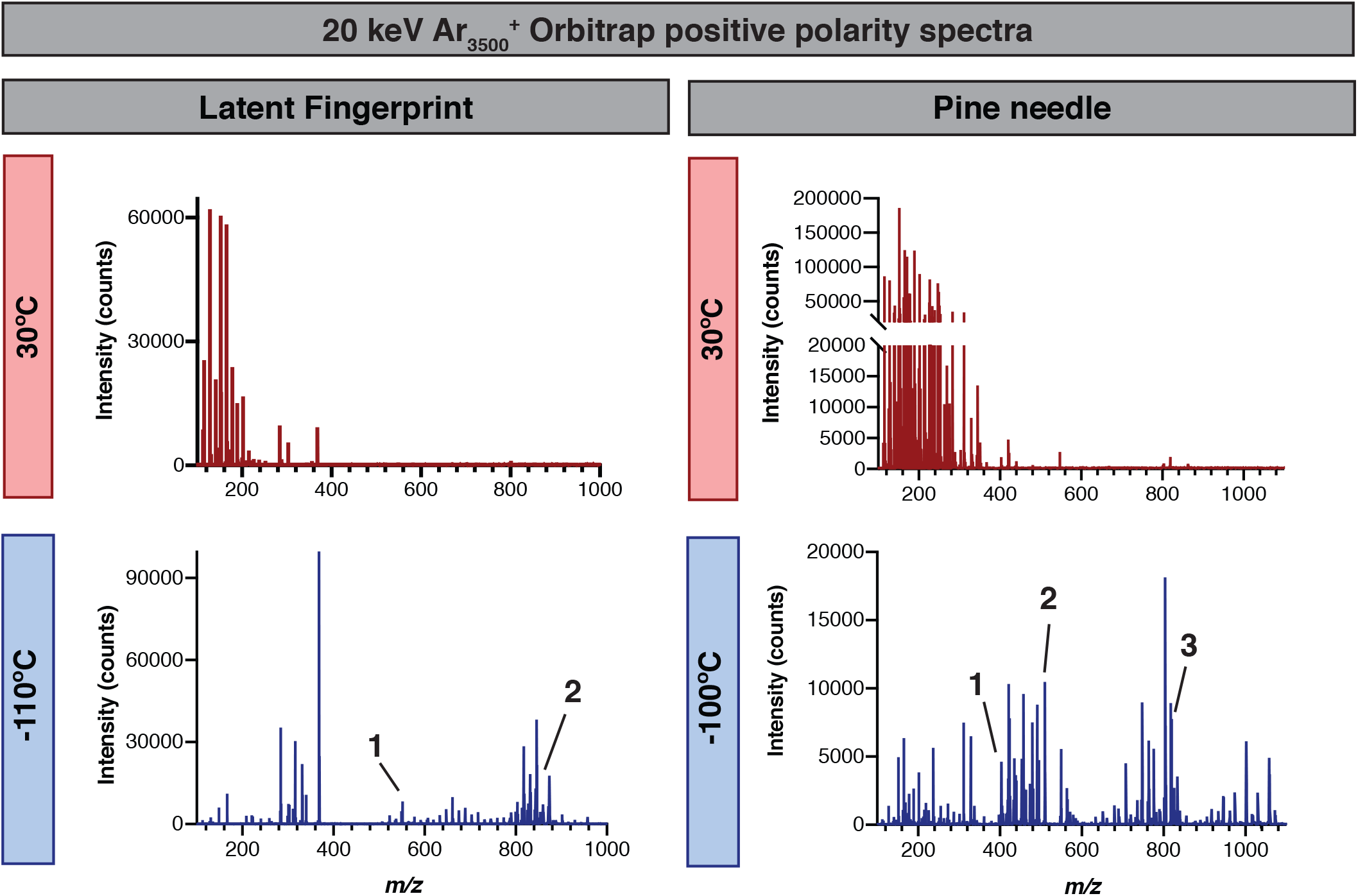
Cryo-OrbiSIMS decreases molecular fragmentation and increases chemical coverage. Comparison of OrbiSIMS spectra of a latent fingerprint and a pine needle with positive polarity using a 20 keV Ar_3500_^+^ GCIB at cryogenic (−100 or −110°C) and ambient (30°C) temperatures. Many more features are detected with cryogenic analysis, including intact triglycerides and semi-volatile hydrocarbons. Fingerprint putative annotations: (1) *m/z* 551.5041, C_35_H_67_O_4_^+^ with mass deviation δ1.4 ppm, C32:0 diglyceride adduct [M+H-H_2_O]^+^ (2) *m/z* 855.741, C_53_H_100_O_6_Na^+^ with mass deviation δ−0.25, C50:1 triglyceride adduct [M+Na]^+^. Pine needle putative annotations: (1) *m/z* 403.4295, C_29_H_55_^+^ with mass deviation δ−0.7 ppm, C29:0 hydrocarbon adduct [M-5H]^+^ (2) *m/z* 509.4561, C_32_H_61_O_4_^+^ with mass deviation δ−0.6 ppm, C29:0 diglyceride adduct [M+H-H_2_O]^+^. (3) *m/z* 819.6379, C_48_H_88_N_2_O_6_P^+^ with mass deviation δ0.6 ppm, C40:5 phosphatidylcholine adduct [M+NH_4_-H_2_O]^+^.

Hydrocarbons, particularly long-chain alkanes, are a major component of the cuticular lipid barrier to water loss in both plants and insects (Blomquist and Bagneres, 2010; Ferveur, 2005; Kunst and Samuels, 2003; Lee and Suh, 2015). We used Cryo-OrbiSIMS to analyze the cuticle on the surface of pine needles of *Pinus nigra* revealing that signal from the ion *m/z* 336.90 provides a specific chemical marker for the guard cells of the stomatal pores (**Fig. 1c**). This contrasts with more spatially uniform epicuticular signals for hexadecenoic acid (*m/z* 255.17) and the volatile hydrocarbon nonacosane (*m/z* 403.09), which we validated with GC-MS (**Supplementary Table 1**). Importantly, these three molecular ions were robustly detected on the surface of pine needles using cryogenic but not ambient temperatures. Analysis of a variety of different plant leaf and fruit samples illustrates the broad diversity of hydrocarbons and other SVOCs that can be detected cryogenically (**Supplementary Table 1**). We also utilized the depth-profiling mode of the Cryo-OrbiSIMS to demonstrate that the semi-volatile hydrocarbon octacosane is restricted to the surface of the cuticle of the gooseberry fruit (**Supplementary Fig. 5a**). The fingerprint and plant cuticle analyses together demonstrate that the new cryogenic workflow expands the chemical space amenable to SIMS imaging to encompass hydrocarbons and other SVOCs, as well as molecules with intermediate vapor pressures like free fatty acids.

The lipid layer that coats the cuticle of the fruit fly *Drosophila melanogaster* is known to be rich in hydrocarbons, wax esters and other lipids (Blomquist and Bagneres, 2010). We set out to test whether cryo-OrbiSIMS could reveal new aspects of the biology of this already well characterized system. Orbitrap spectra of abdominal *Drosophila* cuticle showed a dramatic increase in the number of features that could be detected via cryogenic rather than ambient OrbiSIMS (**Fig 3a, b**). We verified that many of the Cryo-OrbiSIMS peaks correspond to alkane and alkene hydrocarbons, examples of SVOCs that could also be detected in cuticular hexane extracts via GC-MS (**Supplementary Fig. 6**). Utilizing Cryo-OrbiSIMS in imaging mode revealed that the strong signals from hydrocarbons such as tricosene (*m/z* 323.27) and a C_7_H_11_^+^ (*m*/*z* 95.07) fragment are widely distributed across the sternites (hard plates of cuticle) and the pleura (flexible cuticle) (**Fig. 3c**). However, depth profiling shows that tricosene is localized specifically to the surface of the cuticle, known as the epicuticle (**Supplementary Fig. 5b**). Surprisingly, cryo-OrbiSIMS and ambient-OrbiSIMS both reveal that signals from some wax esters are near uniform whereas others are enriched in the pleura or in the sternites (**Fig. 3d**). These non-uniform localizations open up a new and intriguing aspect of insect biology, suggesting that wax esters may define cuticular zones of different barrier properties. Given that we have shown that Cryo-OrbiSIMS is able to detect a wide range of SVOCs and also non-volatile lipids, it is well suited as a screening platform for the metabolic effects of genetic mutations or drugs. As a proof-of-principle, we combined Cryo-OrbiSIMS with spatially resolved genetic manipulations in *Drosophila*. Cells called oenocytes express *Cyp4g1*, a gene encoding a P450 enzyme required for the synthesis of cuticular hydrocarbons (Makki et al., 2014; Qiu et al., 2012). Orbitrap spectra revealed a dramatic decrease in alkanes and alkenes including tricosene following oenocyte-specific knockdown of *Cyp4g1*(*Cyp4g1* RNAi) (**Fig. 3e**). Cryo-OrbiSIMS is therefore a promising new analytical tool for studying the genetic regulation of metabolism with high spatial and chemical resolution.

**Figure 2.**
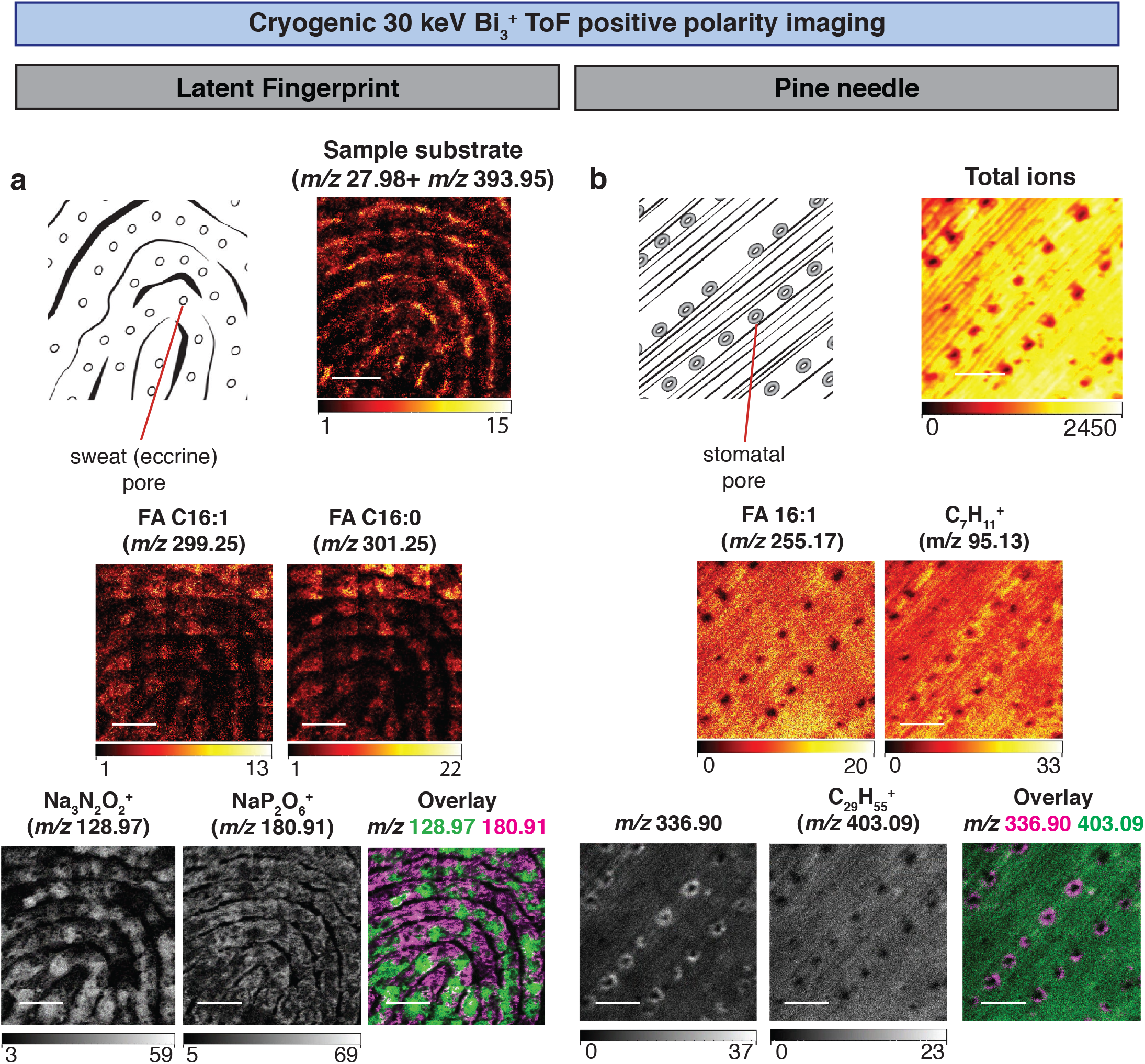
Localization of semi-volatile molecules in pine needles and latent fingerprints using Cryo-OrbiSIMS. a. Positive polarity imaging of a latent fingerprint (diagram indicates eccrine pores) using a 30 keV Bi_3_^+^ LMIG and ToF analyzer. Sample substrate refers to the sum of *m/z* 27.98 (Si^+^) and *m/z* 393.95 (Au_2_^+^), components of the gold-coated silicon wafer. Other mass images show four compounds detected at cryogenic but not at ambient temperatures. Fingerprint putative annotations: *m/z* 299.1956 (detected in Orbitrap and ToF spectra), C_16_H_29_O_2_Na_2_^+^ (Orbitrap mass deviation δ −0.5 ppm), hexadecenoic acid (FA 16:1) adduct [M+2Na-H]^+^. *m/z* 301.2113 (detected in Orbitrap and ToF spectra), C_16_H_31_O_2_Na_2_^+^ (Orbitrap mass deviation δ−0.4 ppm), palmitic acid (FA 16:0) adduct [M+2-Na-H]^+^. *m/z* 128.97, Na_3_N_2_O_2_^+^ localizes close to sweat pores. *m/z* 180.91, NaP_2_O_6_^+^ localizes in a pattern excluding sweat pores. Scale bars represent 0.625 mm. b. Positive polarity imaging of a pine needle (Pinus nigra, diagram indicates stomatal pores) using a 30 keV Bi_3_^+^ LMIG and ToF analyzer. Total ion count image shows position of stomatal pores and m/z 366.90 provides a diagnostic ion for the guard cells surrounding the pores and is detected at cryogenic but not ambient temperatures. Pine needle putative annotations: *m/z* 255.17, C_18_H_35_O_2_^+^, linoleic acid (FA 18:2) adduct [M+H]^+^is detected at both cryogenic and ambient temperatures and with the Orbitrap analyzer (*m/z* 255.2317 and mass deviation δ−0.7 ppm). *m/z* 95.12, C_7_H_11_^+^, likely represents a volatile hydrocarbon fragment and is also detected by EI-GC/MS and OrbiSIMS MS/MS analysis of hydrocarbon standards (**Supplementary Fig. 4**). *m/z* 403.09, C_29_H_55_^+^, C29:0 hydrocarbon adduct [M-5H]^+^ also detected via GC-MS (**Supplementary Table 1**) and with the Orbitrap analyzer (m/z 403.4295 and mass deviation δ−0.7 ppm). Scale bars represent 125 μm. In panels b and c, the intensity ranges (in counts) are represented beneath each mass image.

**Figure 3.**
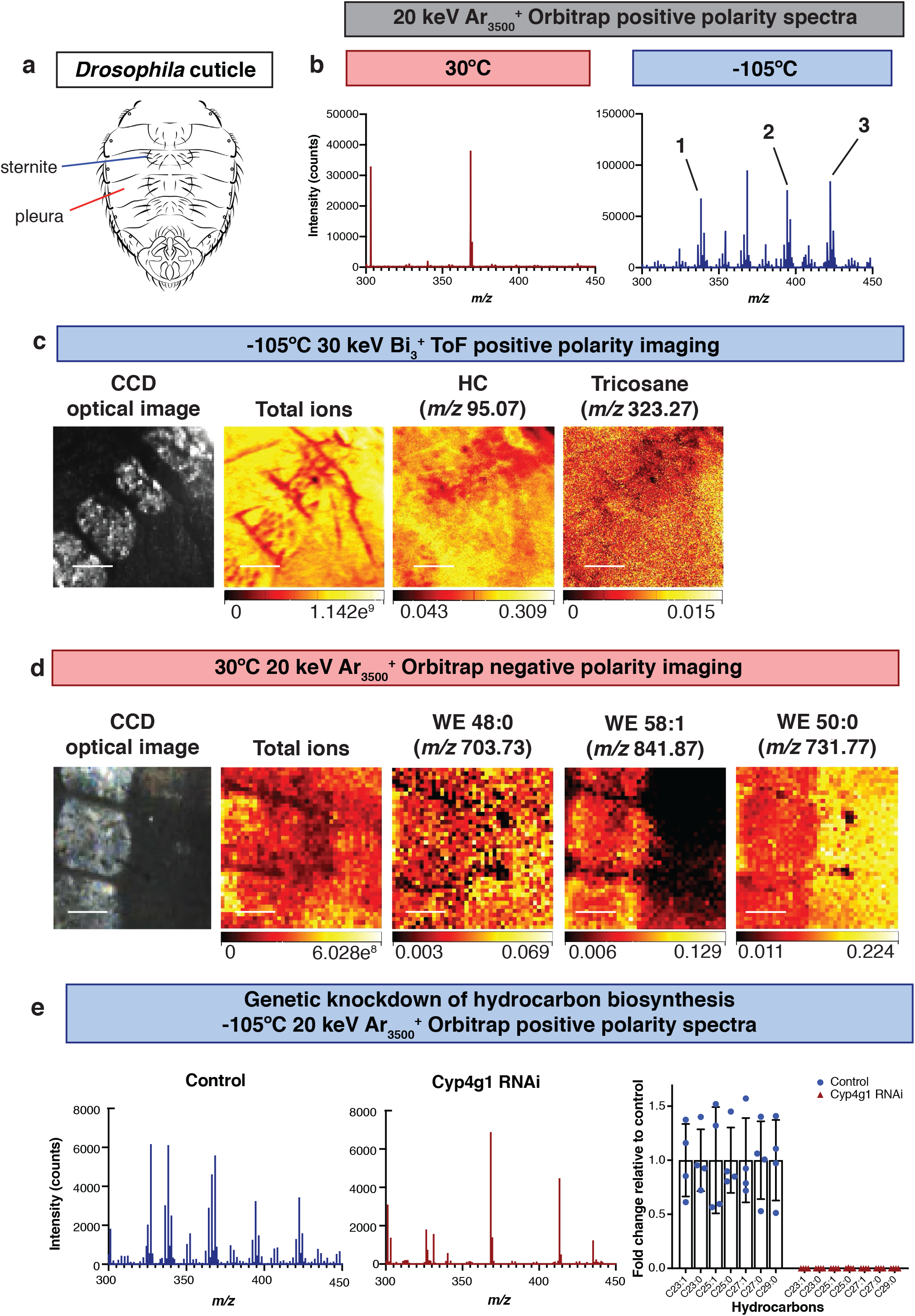
Restricted localization of wax esters and metabolic regulation of hydrocarbons on the Drosophila cuticle. a. The cuticle of the Drosophila abdomen, indicating subdivision into bristle-bearing ventral plates (sternites) and a bristle-free lateral region (pleura). b. OrbiSIMS positive polarity spectra using a 20 keV Ar_3500_^+^ GCIB highlights that many more features are detected at cryogenic (−105°C) than at ambient (30°C) temperatures. Putative annotations: (1) *m/z* 338.378, C_23_H_48_N^+^ with mass deviation δ1.43 ppm, C23:0 hydrocarbon adduct [M+N]^+^, (2) *m/z* 394.440, C_27_H_56_N^+^ with mass deviation δ 0.99 ppm, C27:0 hydrocarbon adduct [M+N]^+^, (3) *m/z* 422.472, C_29_H_60_N^+^ with mass deviation δ1.04 ppm, C29:0 hydrocarbon adduct [M+N]^+^. c. Cryogenic OrbiSIMS ToF imaging in positive polarity using a 30 keV Bi_3_^+^ LMIG. The charge coupled device (CCD) shows the optical image of the acquired area, with sternites and pleura visible and the corresponding total ion counts at each pixel are shown. Abundant cuticular hydrocarbons are widely distributed across the sternites and pleura of the cuticle. Peak annotations: *m/z* 322.2, C_23_H_46_^+^, tricosene, also detected via GC-MS and confirmed with analytical standard. *m/z* 95.07, C_7_H_11_^+^, hydrocarbon (HC) fragment also detected in EI-GC/MS and OrbiSIMS Orbitrap MS/MS spectra of hydrocarbon standards (**Supplementary Fig. 5**). Scale bars represent 125 μm. d. Ambient OrbiSIMS Orbitrap imaging in negative polarity using a 20 keV Ar_3500_^+^ GCIB. The charge coupled device (CCD) shows the optical image of the acquired area, with sternites and pleura visible and corresponding total intensities at each pixel are also shown. Putative annotations: *m/z* 703.7341, C_48_H_95_O_2_^-^ (mass deviation δ 0.5 ppm), C48:0 saturated wax ester (WE 48:0) near-uniform across sternites and pleura. *m/z* 841.8745, C_58_H_113_O_2_^-^ (mass deviation δ −0.1 ppm), C58:1 unsaturated wax ester (WE 58:1) enriched on sternites. *m/z* 731.7656, C_50_H_99_O_2_^-^ (mass deviation δ 0.7 ppm), C50:0 saturated wax ester (WE 50:0) enriched on pleura. Scale bars represent 100 μm. e. Cryo-OrbiSIMS Orbitrap positive polarity spectra using a 20 keV Ar_3500_^+^ GCIB show that cuticular hydrocarbon signals are abundant in genetic control animals (control) but strongly decreased in response to RNAi knockdown of a hydrocarbon biosynthetic enzyme (Cyp4g1 RNAi) - see **Materials and Methods**. Graph shows fold changes of C23 to C29 hydrocarbons in Cyp4g1 RNAi animals versus controls. Bars represent mean, error bars are standard deviation.

Together, the results of this study demonstrate that cryogenic analysis greatly extends the chemical coverage of the OrbiSIMS platform. Combining soft ionization under cryogenic analysis conditions with the high mass resolution of the Orbitrap has revealed unexpected localization patterns for several biological SVOCs. We anticipate that this SIMS advance will have a broad range of applications in metabolic and pharmaceutical research, forensic science and in the analysis of environmental pollutants such as polycyclic aromatic hydrocarbons.

## Materials and Methods

### 3D OrbiSIMS instrumentation and workflow

3D OrbiSIMS (IONTOF GmbH) samples were mounted for cryo-OrbiSIMS analysis (**Supplementary Figure 7**) using a modified a blank Leica VCT holder (Leica microsystems GmbH, cat. #16771610, **Supplementary Figure 8**) to enable the mounting of silicon wafers (or other appropriate substrates) under liquid nitrogen. This modified sample holder is compatible with the 3D OrbiSIMS and forward compatible with IONTOF Hybrid-SIMS (IONTOF GmbH) instruments or other systems that accept Leica EM-VCT holders. After mounting samples under liquid nitrogen in a Leica EM-VCT 500 system (Leica microsystems GmbH), samples were transferred to the instrument loadlock using the Leica EM-VCT 500 cryogenic transfer shuttle at a temperature of about −140 °C and a pressure of 2.0 x 10^-2^ mbar and parked in the loadlock on a cryo-stage cooled by a copper cooling finger until the temperature remained stable at about −135 °C at a pressure of ~1 x 10^-5^ mbar. Samples were then transferred into the analysis chamber and the stage cooled continually during analysis with a mechanically articulated thermal contact allowing movement in the *x*, *y*, *z* plane and cooling to between −100 and −115 °C. Analyses were performed with a 20 keV Ar_3200_^+^ quasi-continuous GCIB analysis beam with a spot size of ~3 μm and a current of 12.6 pA at 15% duty cycle. For depth profiling experiments, the GCIB was operated with a spot size of 20 μm and a current of 0.3 nA at 10% duty cycle. Orbitrap mass calibration was performed using silver clusters generated from a standard. Spectra were acquired using a spot size of 20 μm and represent the sum of 50 scans from an area of 400 x 400 μm and 40 x 40 pixels per scan. Ions were extracted with an extraction field of 2 keV, or 500 V for fly cuticle samples with an extraction delay of 0.925 μs to compensate for sample topography. The Orbitrap was operated in positive-ion polarity with a mass resolution of 240,000 @ 200 *m/z* with an injection time of 2901 ms. Mass spectral information was acquired for the mass range 100 – 1,500 *m/z*. Approximately 144500 shots at 400 μs per cycle were accumulated in the C-trap per acquisition. For MS/MS analysis, a mass window of 0.4 *m/z* was used and the normalized collisional energy set at 1 eV NCE and 45 eV NCE as indicated. To compensate for sample charging, the analysis chamber was flooded with argon gas at a pressure of 1 x 10^-6^ mbar and a flood gun was applied during the analysis with an energy of 20 eV and a current of −10 μA. For ToF imaging experiments, the LMIG was operated using 30 keV Bi_3_^+^ cluster beam in short pulses mode with a current of 0.06 pA. The spectra were calibrated using reference ions of C^+^, CH^+^, CH_2_^+^ and CH_3_^+^. The pine needle ToF image represents a 500 x 500 μm analysis area and 1024 x 1024 pixels per scan in sawtooth raster mode, with the image showing the sum of 35 scans. The total ion dose for this acquisition was 2.75 x 10^9^ ions with 1 shot per pixel and a cycle time of 200 μs. This image was acquired between the temperatures of −112°C and −106°C. The fingerprint ToF image was acquired using a stage scan of a 2.5 x 2.5 cm area and 1250 x 1250 pixels per scan made up of 500 x 500 μm tiles automatically stitched during acquisition, with the image showing the sum of 2 scans. The total ion dose for this acquisition was 2.02 x 10^9^ ions with 1 shot per pixel and a cycle time of 400 μs. The image was acquired between the temperatures of −125°C and −111°C. The *Drosophila* cuticle ToF image represents a 500 x 500 μm analysis area and 1024 x 1024 pixels per scan in sawtooth raster mode, with the image showing the sum of 10 scans. The total ion dose for this image was 7.85 x 10^9^ ions with 1 shot per pixel, with an LMIG current of 0.2 pA and a cycle time of 200 μs. This image was acquired between the temperatures of −101°C and −99°C.

### Data analysis

The 3D OrbiSIMS and ToF SIMS 5 instruments were controlled using SurfaceLab 7.0 (version 7.0.106074, release SL7.0e) and SurfaceLab 6.7 respectively (IONTOF GmbH), with the 3D OrbiSIMS integrating an application programming interface (API) provided by ThermoFisher Scientific(Passarelli et al., 2017). Image and spectral analyses were performed using SurfaceLab 7.0 (IONTOF GmbH), MATLAB 2016a, GraphPad Prism version 8.2.1 and Origin Pro 2018 version SR1. ToF images were processed with 16 pixels binning. *Drosophila* cuticle images (ToF and Orbitrap) were normalized to the total ion count and total intensity respectively to normalize for differences in intensity caused by topography. Chemical structures were drawn using ChemSketch (ACD Labs) version 2018.1.1. For spectral data analysis, a cut off of 5,000 counts was used and a signal-to-noise ratio (SNR) of 1,000. Mass deviation (δ in ppm) was calculated using the average observed mass over three separate measurements. GC-MS data analysis was performed using MassHunter Qualitative Analysis software (Version B.06.00, Agilent Technologies, Inc). Hydrocarbons were identified based on comparisons to spectra and retention times of authentic standards (indicated where available). HESI-Orbitrap data analysis was performed using ThermoFisher XCalibur SP1 Qual Browser version 4.1.50.

All data files associated with this study are available to download from MetaboLights referencing study number MTBLSxxxx.

**See Supplementary Information for Supplementary Figures and the Extended Methods, including sample preparation, TOF-SIMS, GC-MS and HESI-Orbitrap direct infusion methods.**

## Acknowledgements

We acknowledge Alan Ling in Crick Mechanical Engineering for sample holder modifications, Joe Brock and Adrien Franchet for research illustrations, the Crick Media Science Technology Platform, the Crick *Drosophila* facility and Diana Topham for access to plant materials. We thank Gita Mistry and consenting participants for fingerprint samples, Mariana Silva Dos Santos, and the Crick Metabolomics Science Technology Platform for technical advice on HESI-Orbitrap direct infusion, and Alexander Pirkl of IONTOF for technical advice on OrbiSIMS Orbitrap image acquisition. This work was supported by an Investigator Award to APG from the Wellcome Trust (104566/Z/14/Z) and by funding for APG form the Francis Crick Institute, which receives its core funding from Cancer Research UK (FC001088}, the UK Medical Research Council (FC001088) and the Wellcome Trust (FC001088). CLN was supported by an EPSRC iCASE studentship (1938905). This work forms part of the OrbiSIMS project in the Life-Sciences and Health programme of the National Measurement System of the UK Department of Business, Energy and Industrial Strategy.

## References

Blomquist, G.J., and Bagneres, A.-G. (2010). Insect hydrocarbons: biology, biochemistry, and chemical ecology (Cambridge: Cambridge University Press).

Brulet, M., Seyer, A., Edelman, A., Brunelle, A., Fritsch, J., Ollero, M., and Laprévote, O. (2010). Lipid mapping of colonic mucosa by cluster TOF-SIMS imaging and multivariate analysis in cftr knockout mice. Journal of Lipid Research 51, 3034–3045.

Cai, L., Xia, M.C., Wang, Z., Zhao, Y.B., Li, Z., Zhang, S., and Zhang, X. (2017). Chemical Visualization of Sweat Pores in Fingerprints Using GO-Enhanced TOF-SIMS. Anal Chem 89, 8372–8376.

Dickinson, M., Heard, P.J., Barker, J.H.A., Lewis, A.C., Mallard, D., and Allen, G.C. (2006). Dynamic SIMS analysis of cryo-prepared biological and geological specimens. Applied Surface Science 252, 6793–6796.

Ferveur, J.-F. (2005). Cuticular Hydrocarbons: Their Evolution and Roles in Drosophila Pheromonal Communication. Behavior Genetics 35, 279.

Hazarika, P., and Russell, D.A. (2012). Advances in fingerprint analysis. Angew Chem Int Ed Engl 51, 3524–3531.

Ifa, D.R., Manicke, N.E., Dill, A.L., and Cooks, R.G. (2008). Latent fingerprint chemical imaging by mass spectrometry. Science 321, 805.

Kunst, L., and Samuels, A.L. (2003). Biosynthesis and secretion of plant cuticular wax. Prog Lipid Res 42, 51–80.

Kuroda, K., Fujiwara, T., Imai, T., Takama, R., Saito, K., Matsushita, Y., and Fukushima, K. (2013). The cryo-TOF-SIMS/SEM system for the analysis of the chemical distribution in freeze-fixed Cryptomeria japonica wood. 45, 215–219.

Lee, S.B., and Suh, M.C. (2015). Advances in the understanding of cuticular waxes in Arabidopsis thaliana and crop species. Plant Cell Rep 34, 557–572.

Liberles, S.D. (2014). Mammalian pheromones. Annu Rev Physiol 76, 151–175.

Lucattini, L., Poma, G., Covaci, A., de Boer, J., Lamoree, M.H., and Leonards, P.E.G. (2018). A review of semi-volatile organic compounds (SVOCs) in the indoor environment: occurrence in consumer products, indoor air and dust. Chemosphere 201, 466–482.

Makki, R., Cinnamon, E., and Gould, A.P. (2014). The development and functions of oenocytes. Annu Rev Entomol 59, 405–425.

McLafferty, F., and Turecek, F. (1993). Interpretation of Mass Spectra, 4 edn (University Science Books).

Passarelli, M.K., Pirkl, A., Moellers, R., Grinfeld, D., Kollmer, F., Havelund, R., Newman, C.F., Marshall, P.S., Arlinghaus, H., Alexander, M.R., et al. (2017). The 3D OrbiSIMS-label-free metabolic imaging with subcellular lateral resolution and high mass-resolving power. Nat Methods 14, 1175–1183.

Qiu, Y., Tittiger, C., Wicker-Thomas, C., Le Goff, G., Young, S., Wajnberg, E., Fricaux, T., Taquet, N., Blomquist, G.J., and Feyereisen, R. (2012). An insect-specific P450 oxidative decarbonylase for cuticular hydrocarbon biosynthesis. Proc Natl Acad Sci U S A 109, 14858–14863.

Richter, K., Nygren, H., Malmberg, P., and Hagenhoff, B. (2007). Localization of Fatty Acids with Selective Chain Length by Imaging Time-of-Flight Secondary Ion Mass Spectrometry. Microscopy Research and Technique 70, 640–647.

Roussis, S.G., Fitzgerald, W.P., and Cameron, A.S. (1998). Low-energy ionization of hydrocarbons in the quadrupole ion trap mass spectrometer. Rapid Communications in Mass Spectrometry 12, 373–381.

Vrkoslav, V., Muck, A., Cvačka, J., and Svatoš, A. (2010). MALDI Imaging of Neutral Cuticular Lipids in Insects and Plants. Journal of the American Society for Mass Spectrometry 21, 220–231.

Xu, H., and Turlings, T.C.J. (2018). Plant Volatiles as Mate-Finding Cues for Insects. Trends Plant Sci 23, 100–111.

Yew, J.Y., and Chung, H. (2015). Insect pheromones: An overview of function, form, and discovery. Prog Lipid Res 59, 88–105.

Yew, J.Y., Soltwisch, J., Pirkl, A., and Dreisewerd, K. (2011). Direct laser desorption ionization of endogenous and exogenous compounds from insect cuticles: practical and methodologic aspects. J Am Soc Mass Spectrom 22, 1273–1284.

