## Supplementary Information for "Cryogenic OrbiSIMS Localizes Semi-Volatile Molecules in Biological Tissues"

**Antoine Equation:** $$\log_{10}(P) = A - \left(\frac{B}{(T + C)}\right)$$

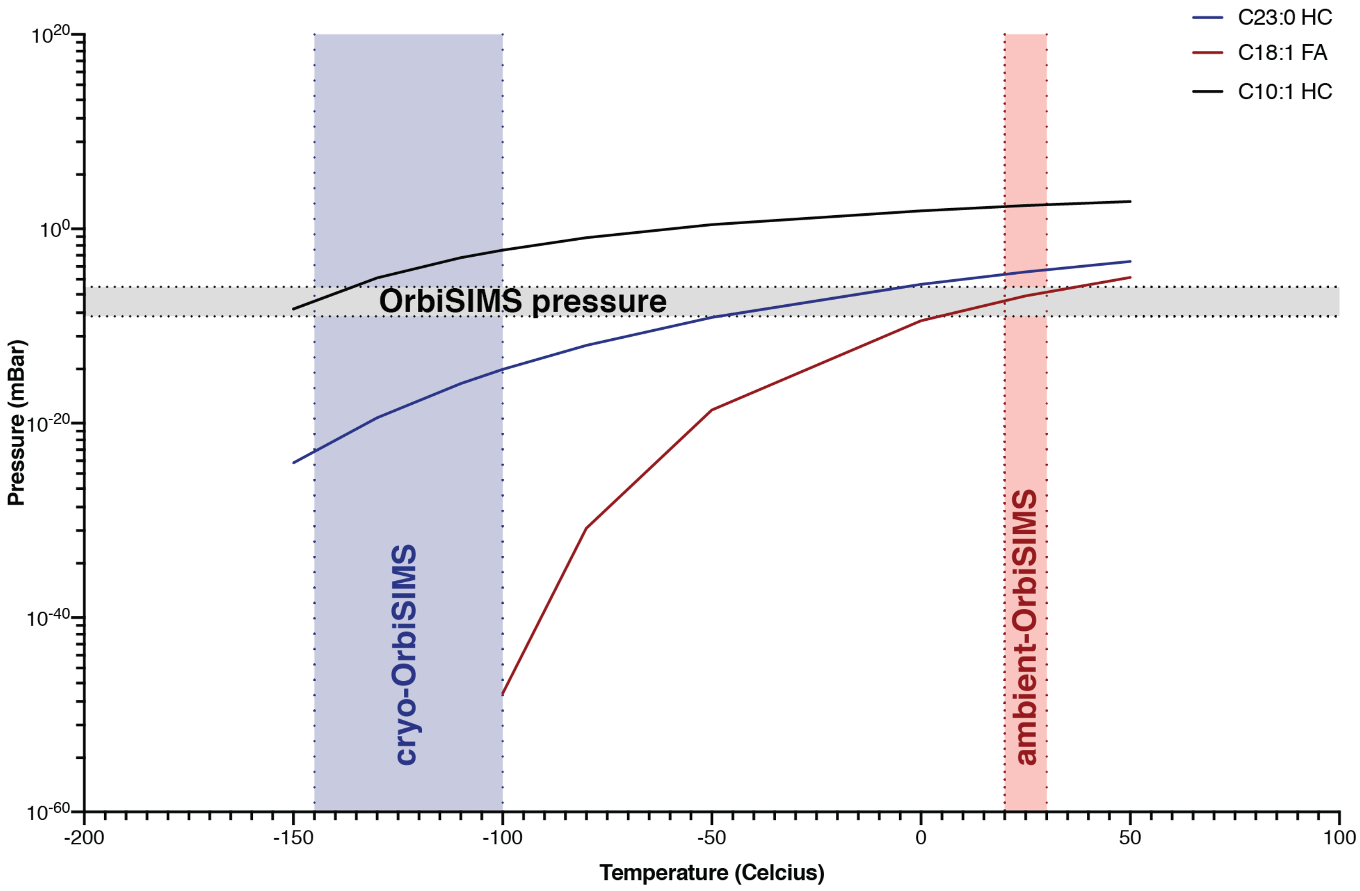


**Supplementary Figure 1: Vapor pressure diagram for semi-volatile lipids**
Curves indicate the predicted pressure and temperature relationship for the vaporisation of two hydrocarbons (C10:1 HC, C23:0 HC) and for oleic acid (C18:1 FA). The standard operating pressure of the OrbiSIMS is indicated at the lower end of the operating range (shaded grey area) with the higher values corresponding to increased pressures using neutral argon gas flooding. The temperature operating ranges of the OrbiSIMS in ambient (red shaded area) and cryogenic (blue shaded area) modes are indicated. Vaporization curves were calculated using the Antoine Equation (indicated) with constants from the NIST database. P = Pressure (bar) and T = Temperature (Kelvin). Constants used: Tricosane A=6.55706, B=4200.069 C=1.864, Oleic acid A=5.04842 B-=2555.604 C=-127.258, and 1-hexene A=3.99063 B=1152.971 C=-47.301. Abbreviations: hydrocarbon (HC), fatty acid (FA).

**a.** **OrbiSIMS with $Ar_{3200}^{+}$ analysis beam**

**9(Z)-Tricosene**
$C_{23}H_{46}$
*m/z* 322.36

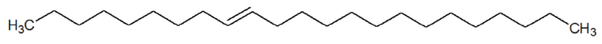

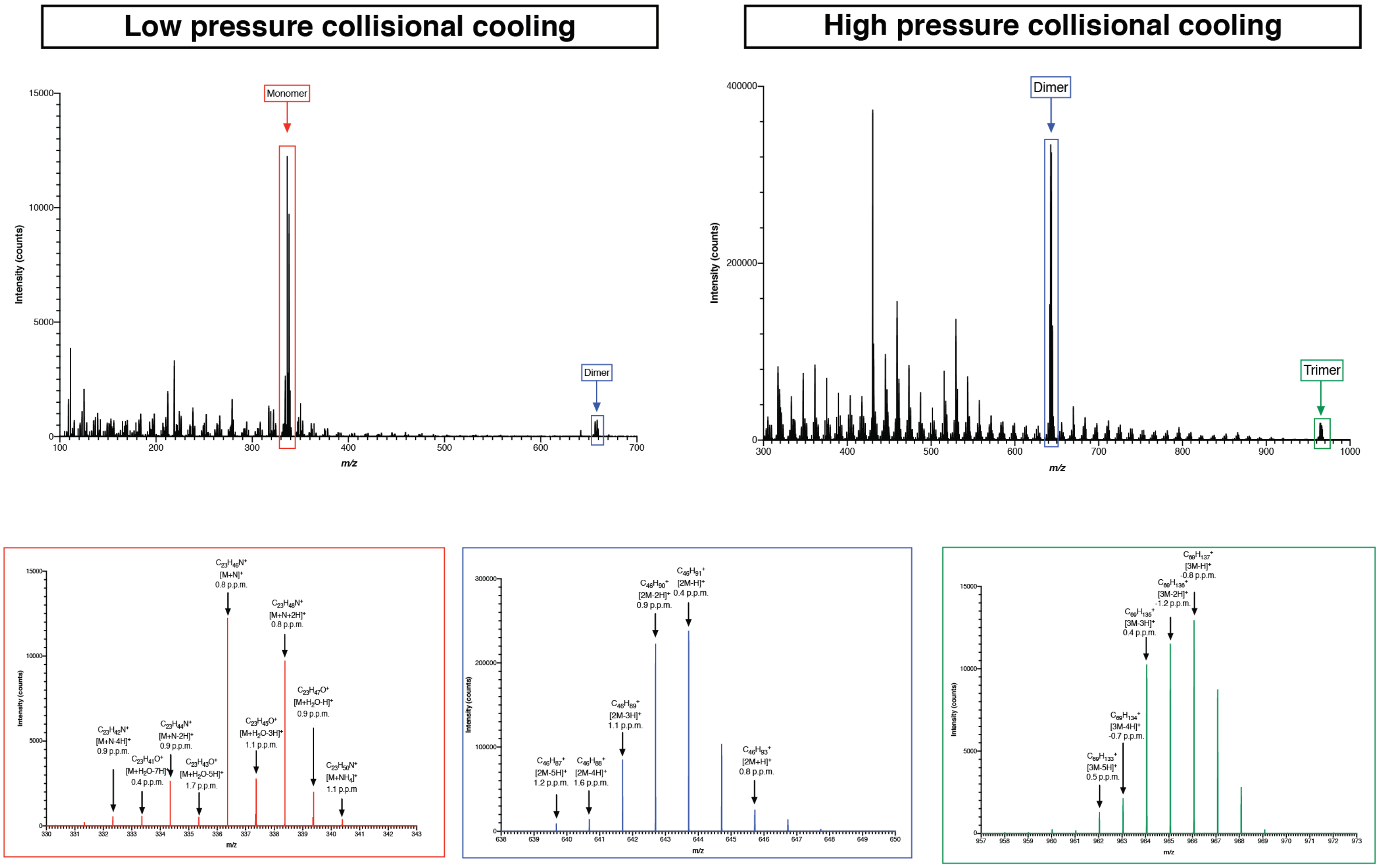


**b.** **ToF-SIMS with 60kV $Bi_3^{++}$ analysis beam**

**7(Z)-Tricosene**
$C_{23}H_{46}$
*m/z* 322.36

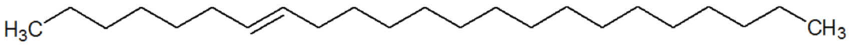

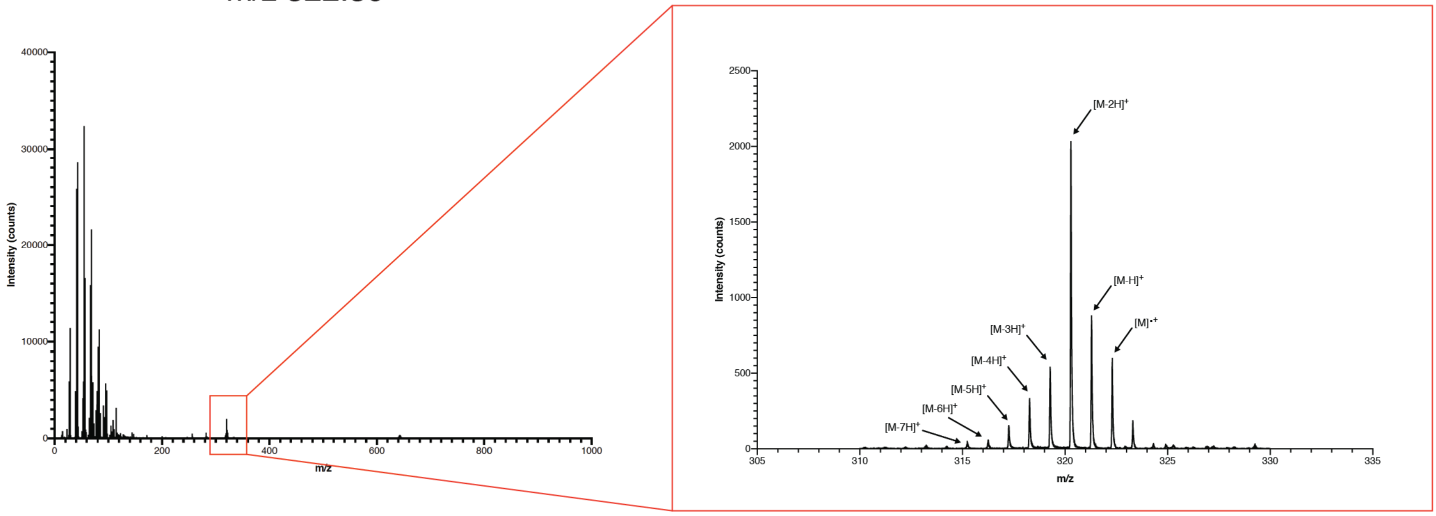

**Supplementary Figure 2: Tricosene adducts**

a. Cryogenic OrbiSIMS GCIB Orbitrap analysis of the hydrocarbon 9(Z)-tricosene using an $Ar_{3200}^{+}$ analysis beam. The most abundant adduct formed incorporates nitrogen, $[M+N]^{+}$. 9(Z)-tricosene also forms dimers and trimers with the predominant adducts likely corresponding to $[2M-H]^{+}$ and $[3M-H]^{+}$. High and low collisional cooling pressures correspond to 1.2 x 10-1 mBar and 4.2 x 10-2 mBar respectively.

b. Cryogenic IONTOF.V LMIG TOF analysis of 7(Z)-tricosene using a $Bi_3^{++}$ analysis beam. Extensive fragmentation is observed, although the molecular ion remains detectable. The predominant adduct likely corresponds to $[M-2H]^{+}$.

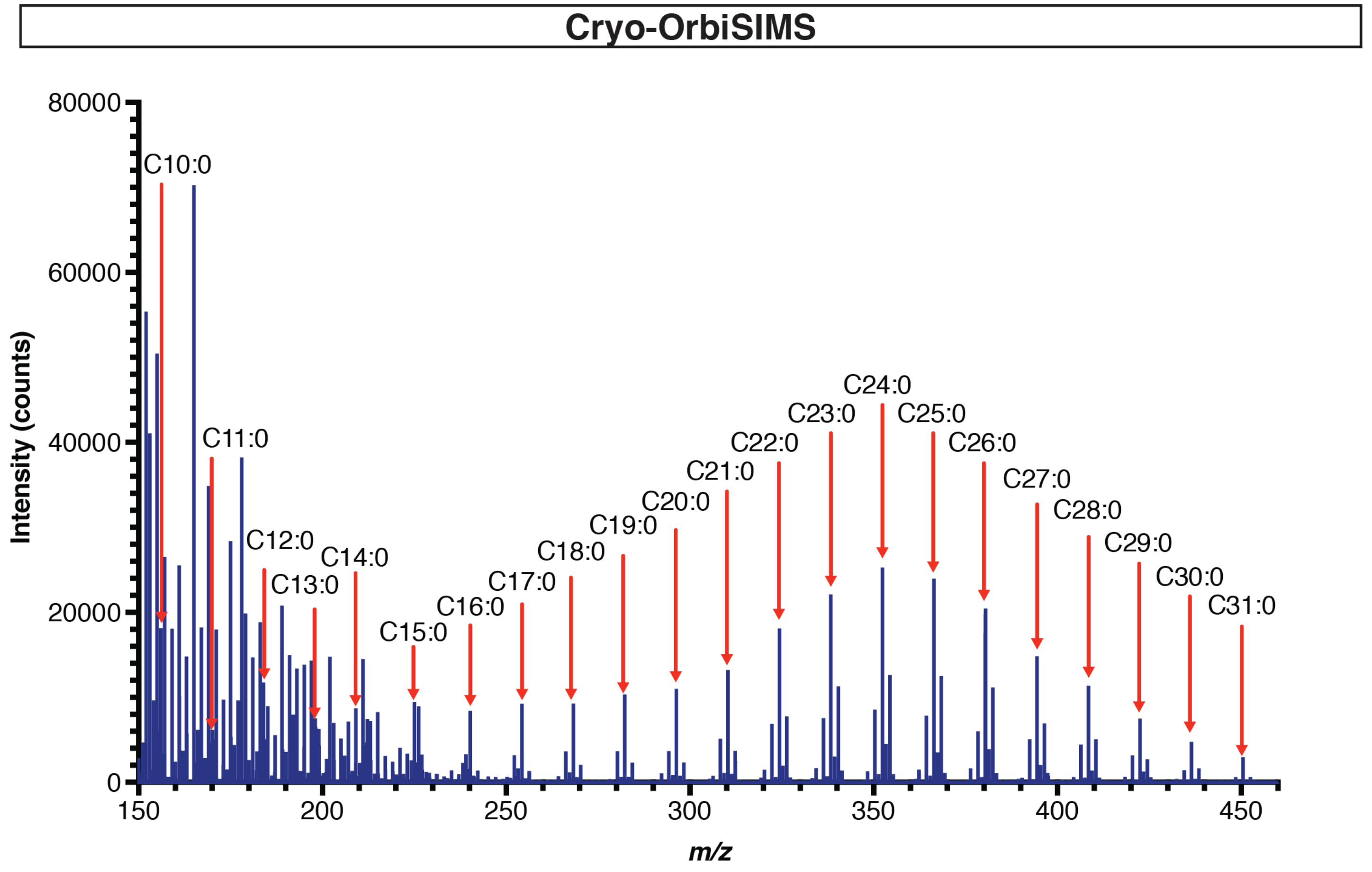


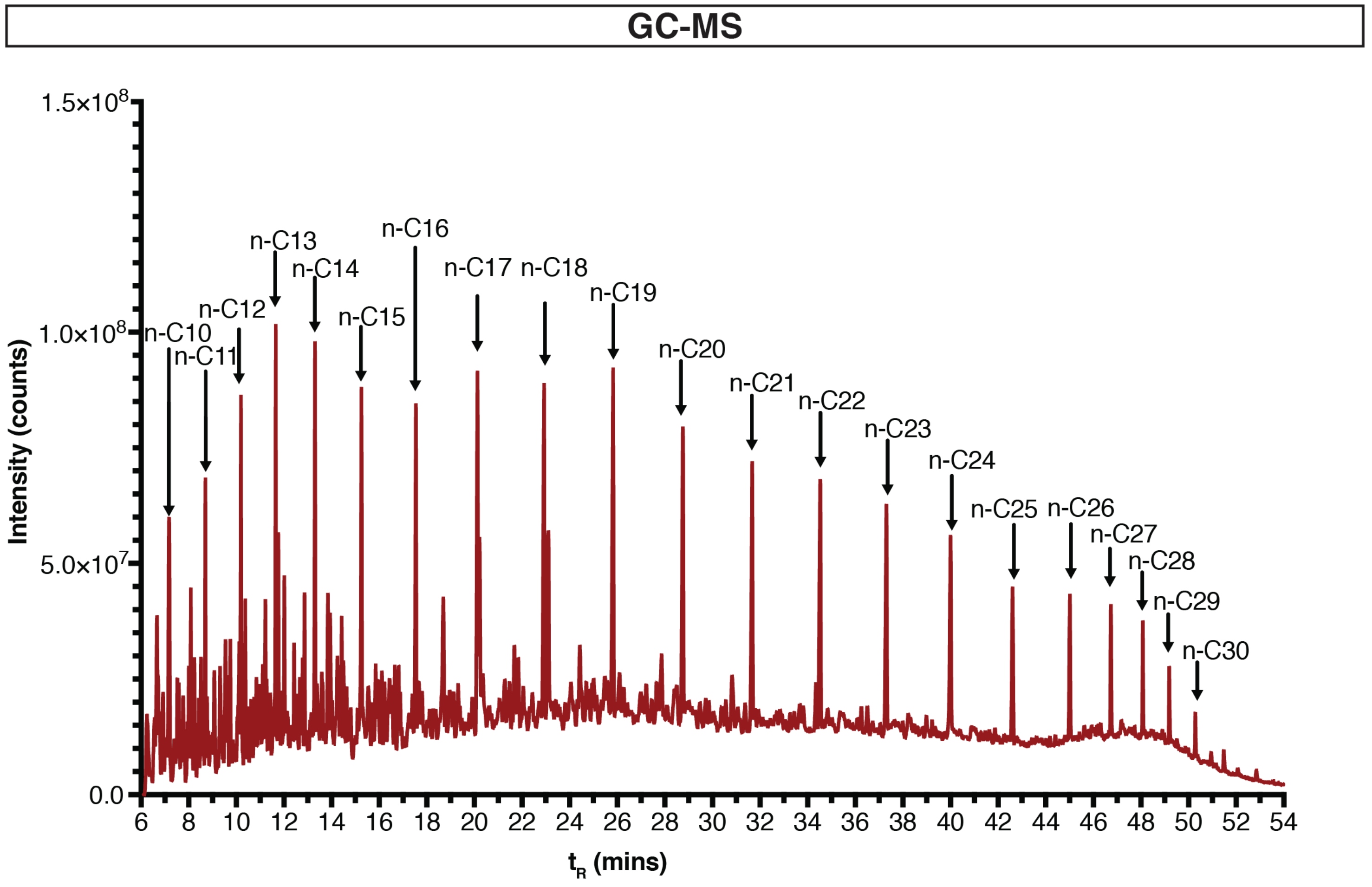


**Supplementary Figure 3: Analysis of gas oil with cryo-OrbiSIMS and GC-MS**
Cryo-OrbiSIMS analysis of ATSM reference gas oil identifies a wide variety of alkanes that are also detected with GC-MS.

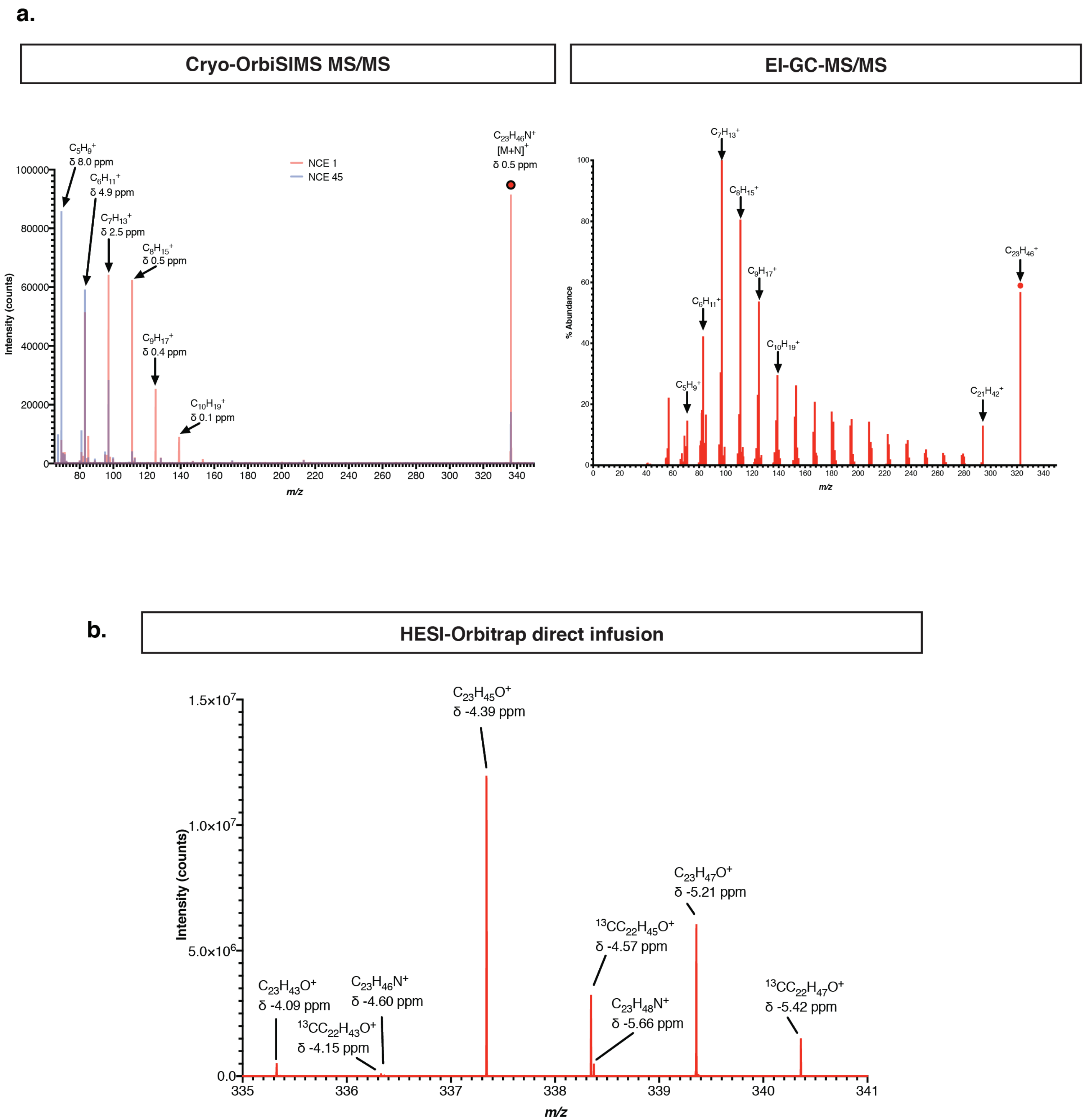


**Supplementary Figure 4: Comparison of Tricosene adducts with three different MS methods.**
a. Cryo-OrbiSIMS MS/MS analysis of the $[M+N]^+$ tricosene adduct reveals fragmentation consistent with a linear hydrocarbon. Increasing the normalized collision energy (NCE) from 1 eV to 45 eV yields complete fragmentation of the parent ion into smaller ions. EI-GC-MS/MS analysis of the $[M]^+$ ion of 9(Z)-tricosene at a NCE of 0.5 eV yields many of the same fragments obtained via Cryo-OrbiSIMS MS/MS.
b. HESI-Orbitrap direct infusion of 9(Z)-tricosene. The most abundant adducts may result from interactions with water but nitrogen adducts, also seen with OrbiSIMS, are visible at lower abundance.

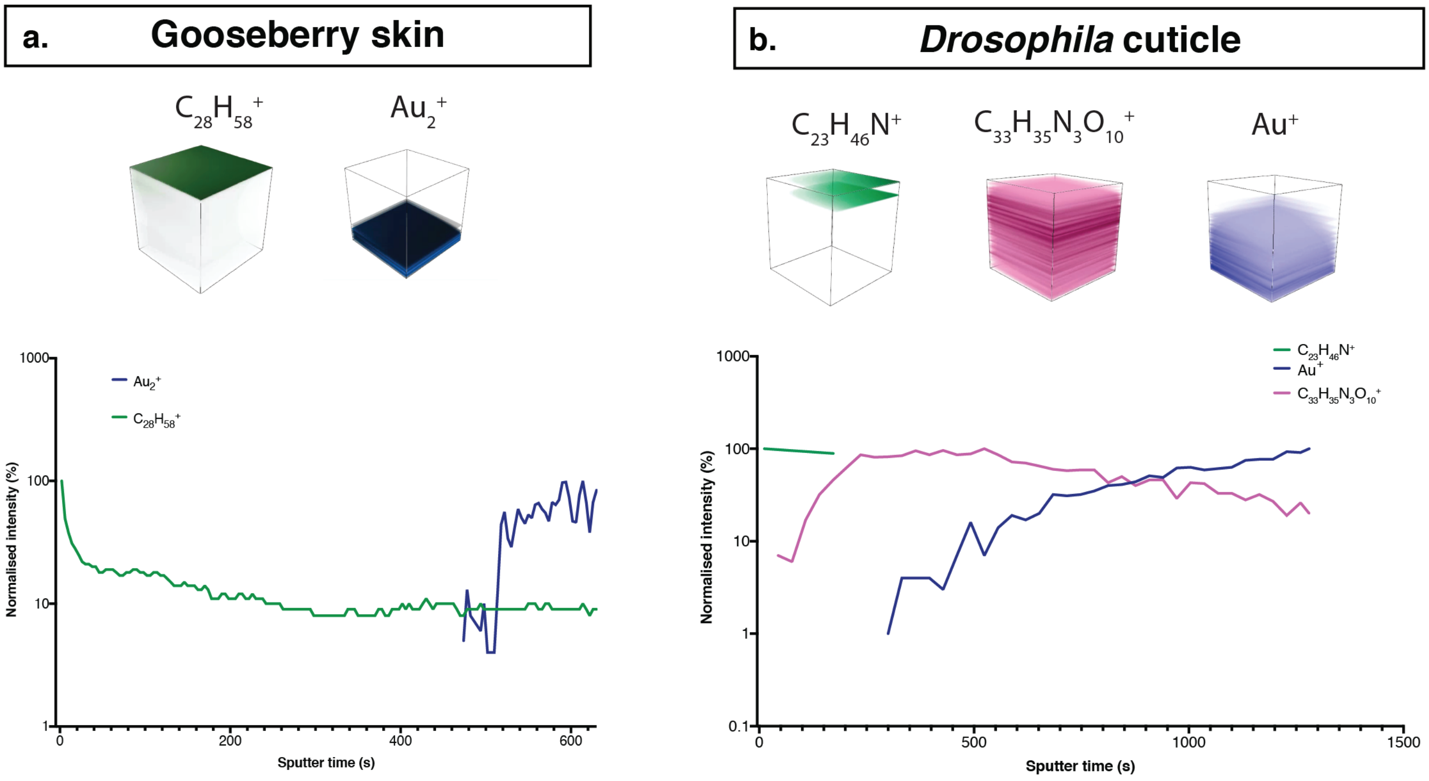


**Supplementary Figure 5: Cryo-OrbiSIMS depth profiling of SVOCs.**
a. Positive polarity depth profiling through the skin of a gooseberry fruit shows that the hydrocarbon octacosane ($C_{28}H_{58}^+$, *m/z* 408.4564, δ -0.1 ppm) is restricted to the surface of the fruit. $Au_2^+$ (*m/z* 393.9326, δ 0.2 ppm) is a reference for the substrate (gold-coated silicon wafer).
b. Positive polarity depth profiling through the Drosophila abdominal cuticle (pleura) shows that tricosene ($C_{23}H_{46}N^+$, *m/z* 336.3628, δ 1.0 ppm) is restricted to the surface of the cuticle. $C_{33}H_{35}N_3O_{10}^+$ (*m/z* 633.2312, δ -0.8 ppm) is a ubiquitous marker of the cuticle, detected throughout the depth profile. $Au^+$ (*m/z* 196.9660, δ -0.2 ppm) is a reference for the substrate (gold-coated silicon wafer). In both panels a and b, the depth profile plot shows signal intensity normalized to 100% for each compound, plotted over sputter time.

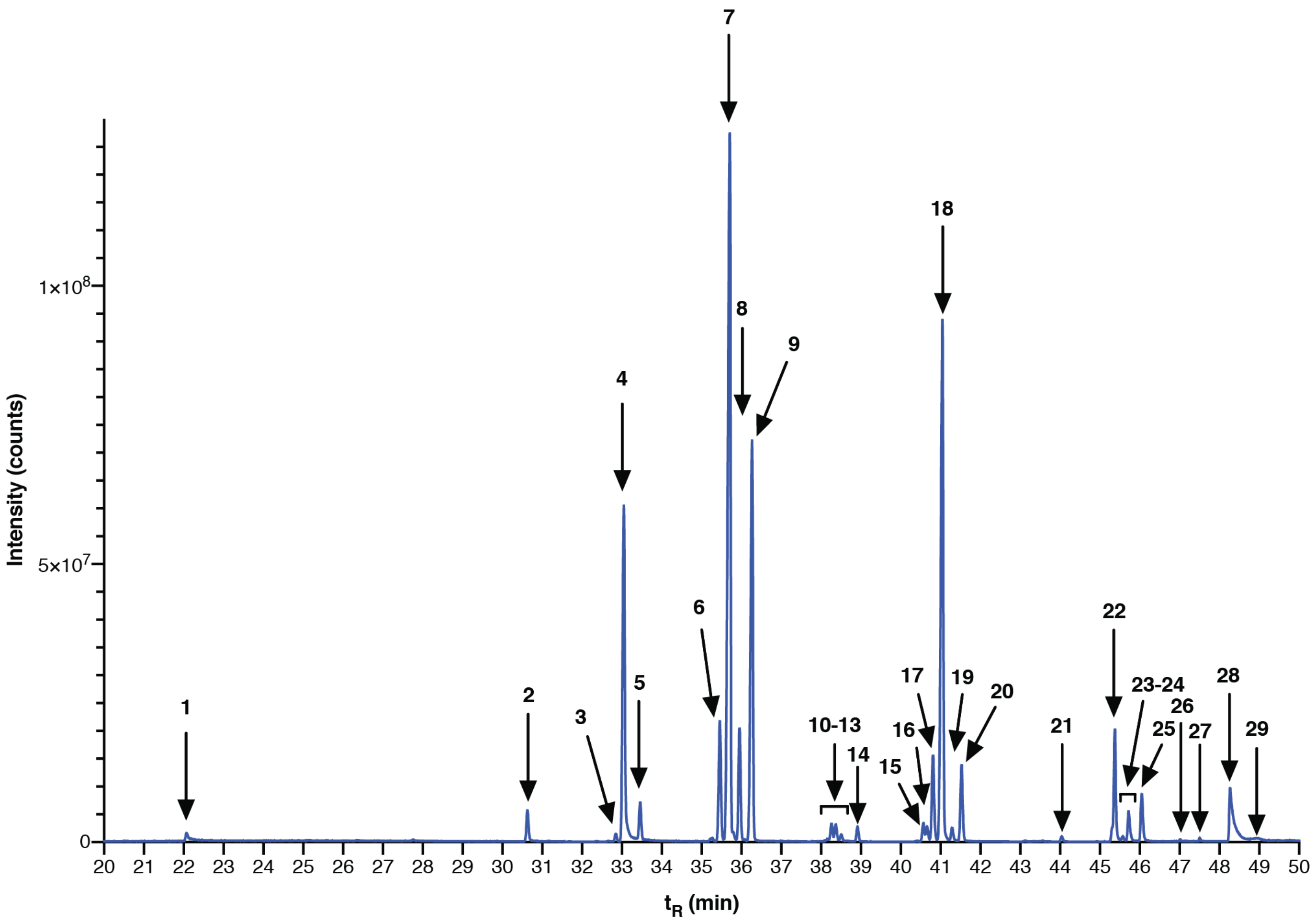


| Peak number | $t_R$ (min) | Parent ion (*m/z*) | Diagnostic ions (*m/z*) | Formula | Putative I.D. | Standard? |
|---|---|---|---|---|---|---|
| 1 | 22.064 | 254.3 | - | $C_{18}H_{38}$ | octadecane | ✓ |
| 2 | 30.626 | 296.3 | - | $C_{21}H_{44}$ | heneicosane | ✓ |
| 3 | 308.2 | 308.2 | - | $C_{22}H_{44}$ | 7-docosene | |
| 4 | 33.049 | 310.2 | 250.1 | $C_{20}H_{38}O_2$ | 11-cis-vaccenyl acetate | ✓ |
| 5 | 33.474 | 310.3 | - | $C_{22}H_{46}$ | docosane | ✓ |
| 6 | 35.455 | 322.3 | - | $C_{23}H_{46}$ | (Z)-9-tricosene | ✓ |
| 7 | 35.705 | 322.3 | - | $C_{23}H_{46}$ | (Z)-7-tricosene | ✓ |
| 8 | 35.946 | 322.3 | - | $C_{23}H_{46}$ | 5-tricosene | |
| 9 | 36.271 | 324.3 | - | $C_{23}H_{48}$ | tricosane | ✓ |
| 10 | 38.165 | 336.3 | - | $C_{24}H_{48}$ | 5-tetracosene | |
| 11 | 38.265 | 336.3 | - | $C_{24}H_{48}$ | 8-tetracosene | |
| 12 | 38.365 | 336.3 | - | $C_{24}H_{48}$ | 7-tetracosene | |
| 13 | 38.506 | 336.3 | - | $C_{24}H_{48}$ | 6-tetracosene | |
| 14 | 38.918 | 338.3 | - | $C_{24}H_{50}$ | tetracosane | ✓ |
| 15 | 40.554 | 352.3 | 337.3, 309.3 | $C_{25}H_{52}$ | x-methyl tetracosane | |
| 16 | 40.663 | 348.3 | - | $C_{25}H_{48}$ | x,x-pentacosadiene | |
| 17 | 40.804 | 350.3 | - | $C_{25}H_{50}$ | 9-pentacosene | |
| 18 | 41.041 | 350.3 | - | $C_{25}H_{50}$ | (Z)-7-pentacosene | ✓ |
| 19 | 41.291 | 350.3 | - | $C_{25}H_{50}$ | 5-pentacosene | |
| 20 | 41.524 | 352.3 | - | $C_{25}H_{52}$ | pentacosane | ✓ |
| 21 | 44.043 | 366.3 | - | $C_{26}H_{54}$ | hexacosane | ✓ |
| 22 | 45.375 | 380.3 | 365.3, 337.3 | $C_{26}H_{54}$ | x-methyl hexacosane | |
| 23 | 45.57 | 378.3 | - | $C_{27}H_{54}$ | 9-heptacosene | |
| 24 | 45.724 | 378.3 | - | $C_{27}H_{54}$ | 7-heptacosene | |
| 25 | 46.049 | 380.3 | - | $C_{27}H_{56}$ | heptacosane | |
| 26 | 47.011 | 394.3 | 351.3 | $C_{28}H_{58}$ | x-methyl heptacosane | |
| 27 | 47.502 | 394.4 | - | $C_{28}H_{58}$ | octacosane | ✓ |
| 28 | 48.268 | 408.3 | 393.3, 365.3 | $C_{29}H_{60}$ | x-methyl octacosane | |
| 29 | 48.955 | 408.3 | - | $C_{29}H_{60}$ | nonacosane | ✓ |

**Supplementary Figure 6: GC-MS validation of Drosophila cuticular hydrocarbons**
GC-MS chromatogram of a hexane extract of the cuticle from five male Drosophila flies, showing the presence of a wide range of different hydrocarbons (as indicated in the table). The majority of these hydrocarbons are also detected by Cryo-OrbiSIMS analysis of Drosophila cuticle.

**a.**

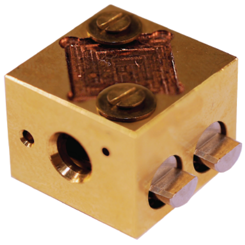

**b.**

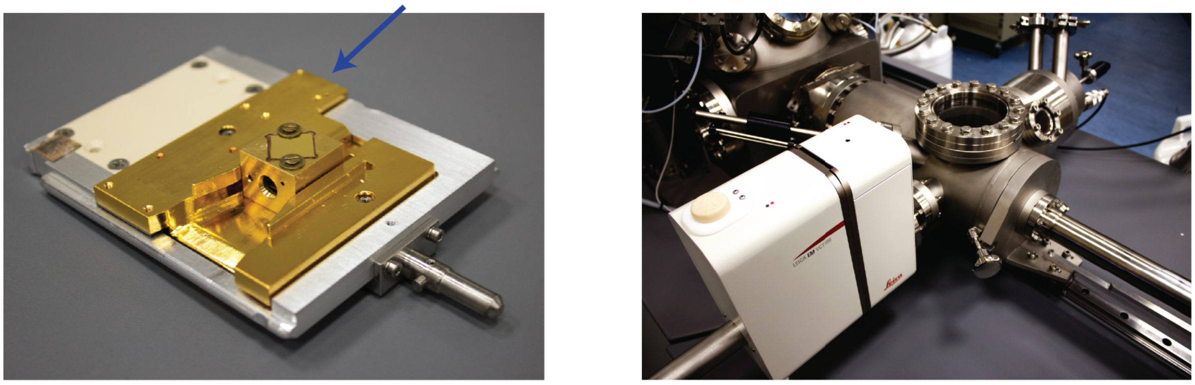

**Supplementary Figure 7: Instrument modifications for Cryo-OrbiSIMS**

a. Modified cryo sample holder. The sample is mounted on a conductive substrate fitting within the 10 mm x 10 mm x 0.5 mm recess. This sample holder is compatible with any SIMS instrument that accepts Leica VCT sample holders.

b. The modified sample holder is transferred under vacuum into the OrbiSIMS via a Leica EM-VCT 500 transfer arm that docks onto the instrument loadlock. The sample holder is then introduced into the IONTOF cryo-stage, which is cooled conductively in situ via a mechanically articulated copper cooling contact as indicated by the arrow.

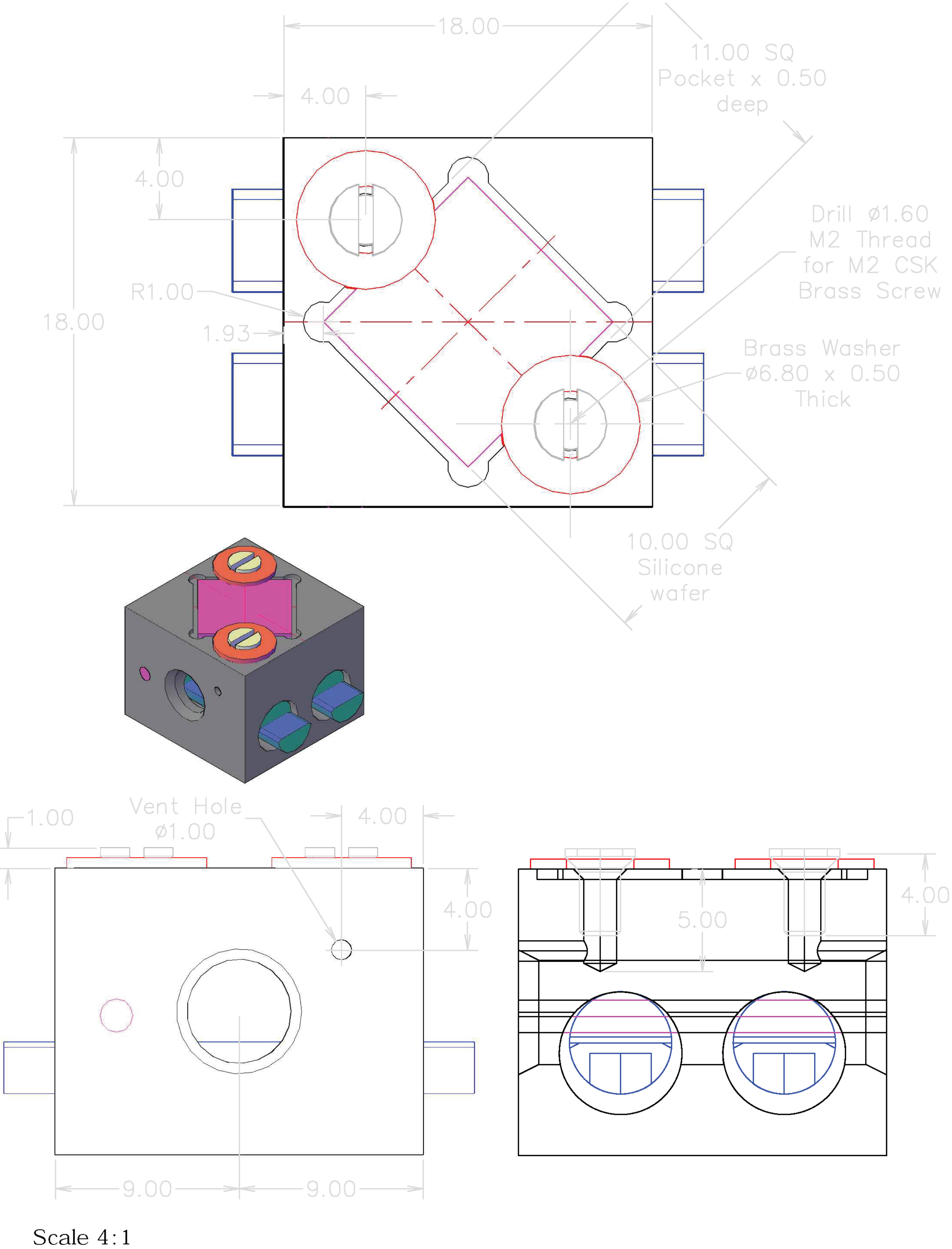


**Supplementary Figure 8:** Leica VCT sample holder (#16771610) modified to accommodate a 10 mm x 10 mm x 0.5 mm silicon wafer by machining a suitably sized depression into a blank sample holder. Screw threads were created and vented orthogonally to allow sample mounting under liquid nitrogen. The sample is held in place using brass washers secured with brass screws.

### Extended Methods

#### Sample preparation

**Plants:** Plants were grown without the use of pesticides or herbicides. The following samples were collected: gooseberry fruit skin (Ribes uva-crispa cv. Hinamaki red), apple tree flower petal (Malus domestica cv. Bountiful), tomato leaf (Solanum lycopersicum cv. Sweet Millions), asparagus skin (Asparagus officinalis cv. Gijnlim), kiwi berry leaf (Actinidia arguta cv. Issai), Victoria plum (Prunus domestica cv. Victoria), Golden Gage plum (Prunus domestica cv. Oullins Golden Gage), and pine needle (Pinus nigra). Samples were collected, mounted on gold coated silicon wafers and stored at 4ºC and analyzed within 24 hours of preparation. The leaf samples were adhered to the gold coated silicon wafer with silver conductive paint (Electrolube, #1LABLSCP/1). For GC-MS, plant samples were washed with 1 mL hexane which was dried in a fume hood. Samples were then resolubilized in 100 µL hexane.

***Drosophila***: All flies were raised on a standard diet composed of on a standard diet composed of 7.5 g agar, 58.5 g glucose, 66.3 g cornmeal, 23.4 g yeast and 19.5 mL of antimycotic solution containing 0.04% Bavistan and 10% Nipagin per litre. For initial experiments, isogenic male *Drosophila* of the w1118 iso31 strain were used. Flies were raised at 25ºC throughout and males separated 24 hours after eclosion and retained at 15 flies per vials until 6 days of age. For oenocyte-specific genetic manipulations, the following strains were used: PromE(800)-GAL4, tub-GAL80TS, UAS-CD8::GFP/Cyo, Dfd::YFP[1] and UAS-Cyp4g1 RNAi[2]. For Cyp4g1 RNAi knockdown, flies were raised at 18ºC during development and then transferred to 29ºC upon eclosion. One of two independent experiments is shown (n=5 samples). A second experiment (n=4 samples) gave similar results. Flies were then separated by sex and genotype after 24 hours then retained at 29ºC at a density of 15 flies per vial until 6 days of age. 6-day old adult male Drosophila melanogaster were used for OrbiSIMS experiments. Flies were anaesthetized with CO2 and dissected in PBS to remove all tissues attached to the cuticle. Dissected cuticles were then washed with deionized water before mounting on cleaned silicon wafers (Agar Scientific # AGG3390-10) coated with gold. Samples were stored at 4ºC, transported on ice and analyzed within 24 hours of preparation. For GC-MS analysis, flies were anaesthetized with CO2 and the surfaces of five flies were extracted with 50 µL hexane containing an internal standard of 0.1mM octadecane for 2 minutes in glass GC vials. The hexane extracts were transferred to glass vials with inserts (Agilent Technologies, Cat No. 5182-0715 and 5183-2085) for immediate analysis. Dimethyl disulfide derivatization was used to determine double bond position for alkenes where no standard was available[3].

**Latent fingerprints:** Fingerprints were collected in accordance with local ethical guidelines from three male and three female consenting volunteers on gold-coated silicon wafers, after a fingerprint conditioning protocol as previously described4. Samples were anonymized and stored in a secure location at 4ºC for 48 hours before analysis.

#### ToF-SIMS

TOF-SIMS 5 (IONTOF GmbH) samples were prepared on ITO coated glass coverslips (SPI supplies #6462-AB 18 mm2, resistivity 70-100 Ω), cut to size with a diamond knife, and mounted on a cooling stage. The loadlock was flooded with nitrogen gas before cooling during pump-down. During the analysis the sample was cooled with a copper cooling finger to ~-110ºC. The analyses were performed using a 60 keV Bi3++ analysis beam with a spot size of ~200nm and a current of 0.025 pA at a cycle time of 200 µs. Secondary ions were extracted with an extraction voltage of 2 keV. Measurements were obtained from a field of view of 500 µm x 500 µm at a resolution of 512 x 512 pixels. Measurements shown are a sum of 20 scans. To compensate for sample charging, a flood gun was applied during analysis with an energy of 21 eV and a current of -10 µA. The total ion dose was ~8.18 x 107 ions/cm$^2$. The mass resolution of the resulting spectra was ~5,000 at m/z 200. Spectral mass calibration was performed in reference to the known ions $C^+$, $CH^+$, $CH2^+$ and $CH3^+$.

**GC-MS**

GC-MS analysis was performed as previously described[1]. In brief, GC–MS was using an Agilent 7890B-5977A GC–MS system in EI mode. Data was acquired and analyzed using MassHunter (Version B.06.00, Agilent Technologies, Inc). Splitless injection (injection temperature 270°C) onto a 30 m + 10 m × 0.25 mm DB-5MS + DG column (J&W, Agilent Technologies) was used, with helium as the carrier gas. The initial oven temperature was 50°C (1 min), followed by temperature gradients to 150°C at 10°C/min, from 150 to 249°C at 3°C/min, from 249 to 300°C at 10°C/min with a hold time of 5 min, and from 300 to 325 °C at 10 °C/min and a hold time of 12 min. Sample running order was randomized using the MassHunter sample sequence randomizer.

**HESI-Orbitrap**

For HESI-Orbitrap direct infusion, 9(Z)-Tricosene was prepared to a concentration of 1mM in hexane. For Orbitrap analysis, the flow rate was 200 µl/min with a nitrogen sheath gas flow rate of 15.54, mass resolving power of 140,000 at m/z 200 and a mass range of 200 – 1500 m/z. The capillary voltage was set at -0.4 V and the capillary temperature at 350 ºC. The spray voltage was set at 4.5 keV.

**References**

1 Stefana, M. I. et al. Developmental diet regulates Drosophila lifespan via lipid autotoxins. Nature Communications 8, 1384, doi:10.1038/s41467-017-01740-9 (2017).

2 Cinnamon, E. et al. Drosophila Spidey/Kar Regulates Oenocyte Growth via PI3-Kinase Signaling. PLOS Genetics 12, e1006154, doi:10.1371/journal.pgen.1006154 (2016).

3 Dunkelblum, E., Tan, S. H. & Silk, P. J. Double-bond location in monounsaturated fatty acids by dimethyl disulfide derivatization and mass spectrometry: Application to analysis of fatty acids in pheromone glands of four lepidoptera. Journal of Chemical Ecology 11, 265-277, doi:10.1007/bf01411414 (1985).

4 Pleik, S. et al. Fatty Acid Structure and Degradation Analysis in Fingerprint Residues. Journal of The American Society for Mass Spectrometry 27, 1565-1574, doi:10.1007/s13361-016-1429-6 (2016).
